# A chemical genomic screen identifies novel genetic interactors of the methylation-dependent *de novo pathway* of phosphatidylcholine biosynthesis in *Saccharomyces cerevisiae*

**DOI:** 10.1101/2025.01.07.631756

**Authors:** Amanda Maliva, Wayne R. Riekhof

## Abstract

Phosphatidylcholine (PtdCho) is the most abundant phospholipid in most eukaryotic cell and organelle membranes, and cells must regulate its synthesis and interorganelle transport to maintain correct membrane compositions and biophysical properties. Our knowledge of genes encoding enzymes of PtdCho biosynthesis is largely complete, and a great deal is understood about their localization and regulation, however our understanding of molecular mechanisms regulating PtdCho biosynthesis and trafficking remains incomplete. To identify genes that show epistatic relationships with the methylation pathway of PtdCho biosynthesis, we performed a chemical-genomic screen of a *Saccharomyces cerevisiae* gene-deletion mutant collection using a phosphatidylethanolamine (PtdEtn) methyltransferase inhibitor, 2-hydroxyethylhydrazine (HEH). HEH functions by selectively inhibiting the PtdEtn methyltransferase enzymes Pem1p/Cho2p and Pem2p/Opi3p. We demonstrate that the addition of exogenous choline or lyso-phosphatidylcholine can recover HEH-mediated growth inhibition, and used this finding to design a functional-genomic screen to identify genes which, when deleted, render the strain unable to grow when the methylation pathway is partially inhibited. We now report the identification of 410 *S. cerevisiae* gene deletion mutants that exhibit HEH hypersensitivity, and identify among those a core set of 21 genes that are known to epistatically interact with genes encoding enzymes of the PtdEtn methylation pathway. This gene set was enriched in functions relating to glycerolipid and sterol biosynthesis and their regulation, the high-osmolarity glycerol (HOG) pathway, and genes involved in chromatin remodeling and transcriptional regulation. These results demonstrate that PtdCho produced by any one of the Kennedy pathway, methylation pathway, or acyltransferase pathway can maintain necessary cellular PtdCho compositions, but that disruption of any one of these pathways leads to epistatic interactions with non-overlapping subsets of genes, thus providing new insights on the specific functions of these pathways. The design and implementation of this screening strategy establishes HEH as a useful tool for specific inhibition of the methylation pathway in high-throughput functional-gnomic screens, which will facilitate further studies on the synthesis, transport, and function of PtdCho.

## Introduction

*Saccharomyces cerevisiae* makes use of three independent pathways for the synthesis of the major membrane lipid, phosphatidylcholine (PtdCho; **See Fig. 1B & 7**). In this work, we refer to these as the methylation (M), Kennedy (K), and acyltransferase (A) pathways. PtdCho from the M-pathway is produced by *N*-trimethylation of phosphatidylethanolamine (PtdEtn), catalyzed by the sequential action of the methyltransferases Cho2p/Pem1p and Opi3p/Pem2p (1, 2). The K-pathway converts free choline (Cho) to PtdCho using cytidinediphosphate-choline (CDP-Cho) as a central intermediate, and requires the activity of Cki1p (Cho kinase, generating phosphocholine), Cct1p (phosphocholine : CTP cytidylyltransferase), and Cpt1p (diacylglycerol : CDP-Cho cholinephosphotransferase) (3, 4). Lastly, lyso-phosphatidylcholine (1-*O*-acyl-2-hydroxyl-3-glycerophosphocholine; lyso-PtdCho) can be taken up and converted to PtdCho via the Ale1p acyltransferase-dependent exogenous lyso-lipid metabolism pathway (A-pathway) (5, 6). PtdCho is the end-product of each of these three pathways, and in *S. cerevisiae,* PtdCho produced from any of these pathways in isolation is sufficient to support membrane biogenesis and cell growth and division under standard laboratory conditions (7). However, while PtdCho produced by any one of these pathways in the absence of the others is sufficient for viability, it is also clear that these three biosynthetically distinct pools of PtdCho are not entirely functionally equivalent. Specifically, the diacylglycerol moieties of PtdCho produced via the methylation and Kennedy pathways are distinct, with more unsaturated molecular species being produced via the M-pathway relative to the K-pathway (1,8). The cellular functions of these pathways are also distinct, since defects in the K-pathway, but not M- or L-pathways, were isolated as bypass suppressors of the essential functions of the Sec14p lipid-binding protein (9–11), and PtdCho derived from the K-pathway is specifically and preferentially degraded by the lipase Nte1p upon heat shock (12). This situation is analogous to the synthesis of the other major aminoglycerophospholipid, PtdEtn, in which PtdEtn pools synthesized by different biosynthetic pathways are functionally distinct in their ability to support mitochondrial function (13–15). These observations indicate that specific, biosynthetically distinct and differentially localized pools of PtdEtn and PtdCho in the yeast cell are used for specific and non-overlapping cell biological functions.

**Figure 1:**
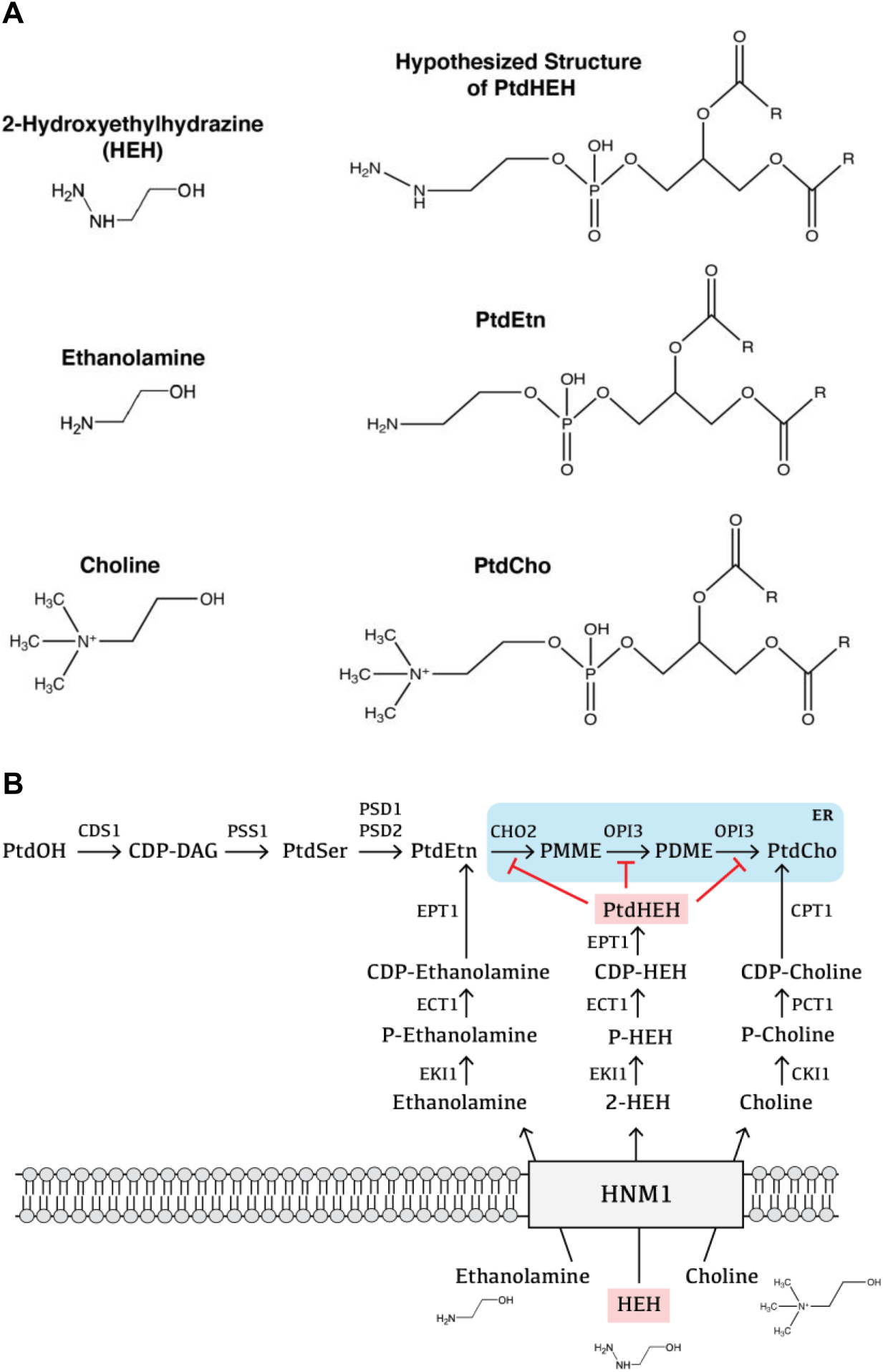
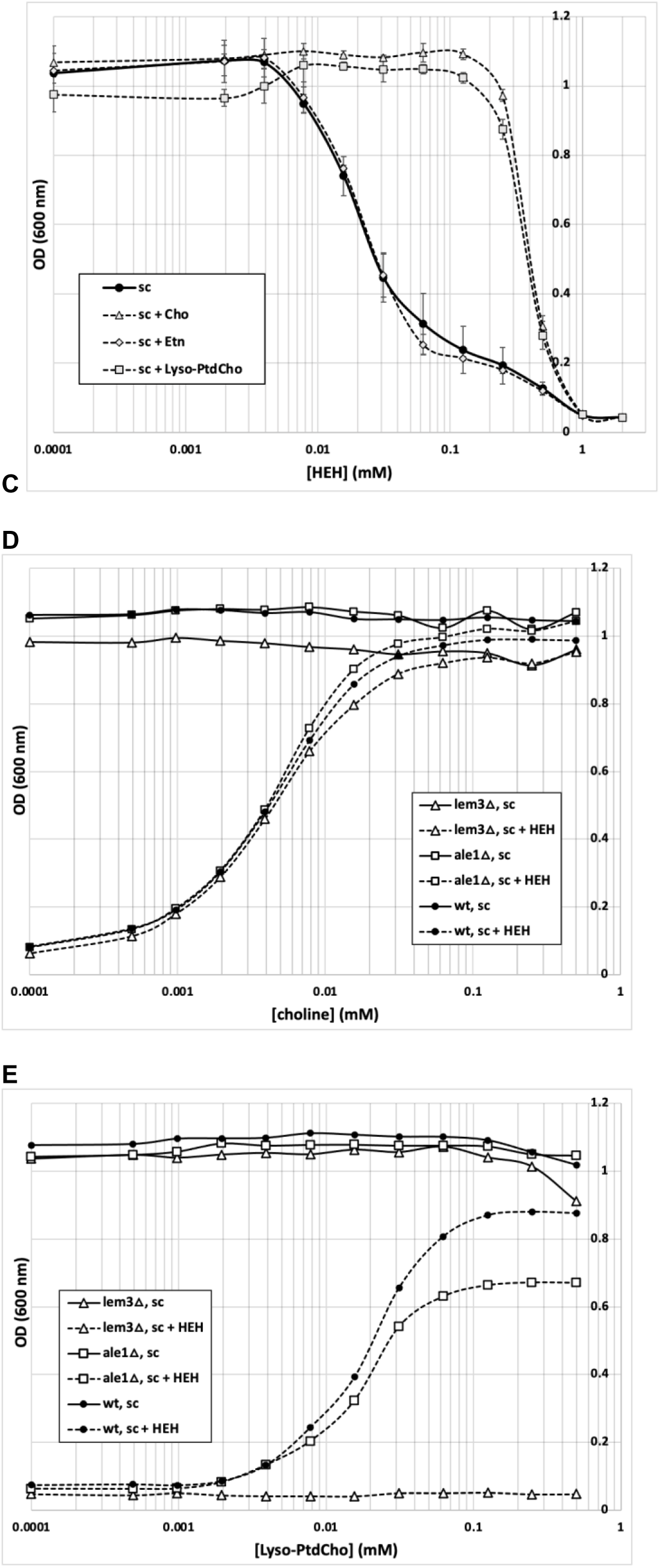
2-Hydroxyethylhydrazine (HEH) is a PtdCho methylation pathway inhibitor. *A*, chemical structures of the methylation pathway derivatives & phospholipids, 2-Hydroxyethylhydrazine (HEH) and the theorized HEH-based phospholipid inhibitor. *B*, pathway diagram of hypothesized HEH-mediated disruption of the methylation pathway. *C*, wildtype growth sensitivity to HEH in synthetic complete liquid media following choline (Cho), ethanolamine (Etn), or lyso-phosphatidylcholine (Lyso-PtdCho) supplementation. *D*, choline-mediated growth recovery of *ale1Δ*, *lem3Δ*, and WT following supplementation of HEH. *E*, Lyso-PtdCho-mediated growth recovery of *ale1Δ*, *lem3Δ*, and WT following supplementation of HEH.

As a structural component of yeast membranes, PtdCho is the dominant species, typically constituting ∼50 mol% of glycerophospholipids (4). The synthesis and interorganelle transport of PtdCho is, under most circumstances, an essential function, although PtdCho synthesis can be rendered dispensable under certain scenarios, such as when the growth of a Cho-auxotrophic *cho2/pem1*Δ *opi3/pem2*Δ double mutant strain is supported by the provision of propanolamine, leading to the synthesis of phosphatidylpropanolamine in place of PtdCho (16, 17). We have also recently shown that PtdCho can be replaced by the phosphorus-free betaine lipid diacylglyceryl-*N,N,N*-trimethylhomoserine when a fungal betaine lipid synthase (Bta1p) is expressed in a *cho2/pem1*Δ *opi3/pem2*Δ strain, which effects reversion of this strain to Cho prototrophy in the absence of all PtdCho synthesis pathways (18). Given this flexibility and apparent redundancy in yeast membrane biogenesis, genetic screens for the identification of new genes involved in lipid biosynthetic and transport pathways are often conducted in strains in which one or more of these pathways have been rendered non-functional through the introduction of inactivating mutations (4). Our current report details a new approach to identify PtdCho biosynthesis and transport mutants, using chemical inhibition of specific PtdCho biosynthetic activities to screen for mutations that render the remaining pathway(s) insufficient to support cell growth.

To begin to genetically dissect the specific cellular functions of different biosynthetic pools of PtdCho derived from the M, K, and L-pathways, we designed a mutant screen to identify genes which, when deleted, render cells unable to grow using only the K- or L-pathways of PtdCho synthesis to support membrane biogenesis and cell growth under respiratory conditions. The basis for our screen (**Fig. 1B**) stems from the observation that 2-hydroxyethylhydrazine (HEH) can act as a specific inhibitor of PtdEtn methylation (M-pathway) (19), thus chemically recapitulating the Cho auxotrophy of a *cho2/pem1*Δ *opi3/pem2*Δ double deletion strain. HEH is taken up via the Hnm1p transporter, through which Cho and the structurally related nitrogen mustard mechlorethamine are transported into the cell (19, 20). After uptake, HEH is presumed to enter the Kennedy pathway to form phosphatidyl-2-hydroxyethylhydrazine (PtdHEH), which acts as an irreversible inhibitor of the Cho2p/Pem1p and Opi3p/Pem2p methyltransferases, leading to cell death in the absence of Cho or lyso-PtdCho (19). We now report the screening and identification of HEH-hypersensitive mutants in a haploid whole-genome deletion collection by culturing the arrayed mutants on solid medium containing a concentration of HEH (400 µM) that is tolerated by the wild type. Thus, the mutants we identify are unable to grow when the methylation pathway is partially inhibited. The ability of exogenously provided Cho or lyso-PtdCho to support growth under these conditions was further assessed, thus defining the optimal concentrations of these amendments and their ability to support the growth of the hypersensitive mutants.

## Results

### 2-Hydroxyethylhydrazine is a PtdCho methylation pathway inhibitor, chemically inducing choline or lyso-PtdCho auxotrophy

An earlier study demonstrating the use of 2-hydroxyethylhydrazine (HEH) as an inhibitor of the *N*-trimethylation of PtdEtn to form PtdCho (19) prompted us to explore its utility for studying the methylation pathway (M-pathway) of PtdCho biosynthesis using a chemical-genomics screening approach. **Figure 1B** shows the overall scheme, in which HEH enters the cell via the Hnm1p choline transporter, and subsequent metabolism by Kennedy pathway enzymes results in the formation of phosphatidyl-2-hydroxyethylhydrazine (PtdHEH), which was proposed as an irreversible inhibitor of the PtdCho methyltransferases Pem1p/Cho2p and Pem2p/Opi3p (19). As illustrated in **Figure 1A**, HEH differs in structure to Etn by the presence of a hydrazine function, rather than a primary amine group. This structural similarity is the basis for the HEH uptake by the Hnm1p transporter, which has a promiscuous substrate specificity, and can transport Etn, Cho, and other small amines, such as the nitrogen mustard mechlorethamine, the metabolic cofactor carnitine, and the osmoprotectant glycine betaine. As described above, subsequent metabolism via the Kennedy pathway is proposed to convert HEH into a form that acts as a PtdCho methyltransferase-specific inhibitor (**Fig. 1B**).

To validate that these results are a specific consequence of disrupted PtdEtn methylation, the growth of the WT strain (BY4741) in minimal media (SC Glc) was observed with increasing concentrations of HEH. Cultures were supplemented with 0.5 mM Cho, Etn, or lyso-PtdCho to determine the capacity of these exogenous supplements to support growth when the methylation pathway is inhibited by HEH. **Fig. 1C** shows that optimum growth of the WT strain is compromised at a concentration of HEH above ∼8 µM, with 50% growth inhibition (IC_50_) at ∼100 µM, and complete inhibition at 0.5 to 1 mM HEH. The addition of Etn failed to rescue HEH-inhibited growth and yielded a growth vs. concentration curve essentially identical to the unsupplemented condition. Because the PtdCho M-pathway, i.e. *N*-trimethylation of PtdEtn, is the principal target of HEH, the provision of exogenous Etn was not expected to recover HEH inhibited growth. However, this result shows that Etn does not suppress the cytotoxicity of HEH by competition for the Hnm1p transporter. Conversely, addition of either 0.5 mM Cho or lyso-PtdCho restored growth of HEH-inhibited cultures in SC Glc, increasing the apparent IC_50_ more than 10-fold. In contrast to Etn, addition of Cho or lyso-PtdCho both supply the alternate pathways of PtdCho formation by the Kennedy (K-pathway) or acyltransferase (A-pathway) pathways, respectively, thus alleviating the growth inhibition due to loss of PtdEtn-derived PtdCho biosynthesis. Taken together, these results demonstrate that HEH inhibition of the M-pathway is the principal mechanism of cell growth inhibition in liquid media at concentrations between ∼10-250 µM, and that at concentrations above this threshold, growth inhibition by HEH is likely a non-specific effect of its reactive hydrazine functional group, independent of its effects on PtdCho biosynthesis.

We next analyzed the lipid profile of the WT strain when subjected to increasing concentrations of HEH (0, 0.15, and 0.5 mM HEH). Thin layer chromatography revealed a dramatic decrease in both PtdCho and PtdEtn formation (**Fig. 2A**). We also observed the formation of a novel lipid species in the HEH-treated samples, which was found to increase in abundance with increasing HEH concentration. The absence of this band in the untreated control led us to suspect its identity as the inhibitory PtdHEH compound. Repeated attempts to confirm the identity of this species using LC-MS and NMR were unsuccessful and gave conflicting results, and it seems this species may be unstable, and is labile in the conditions used for sample preparation after TLC separation, as might be expected for an organic hydrazine. Additional analyses of HEH-treated and -untreated WT and *cho2Δ/pem1Δ* cultures, with authentic PMME (Phosphatidylmonomethylethanolamine) and PDME (Phosphatidyldimethylethanolamine) lipid standards, additionally confirmed that the novel lipid is not a PtdCho methylation pathway derivative (**Fig. 2C**), however the novel lipid ultimately remains unidentified, and further analyses to confirm its identity are beyond the scope of this work.

**Figure 2:**
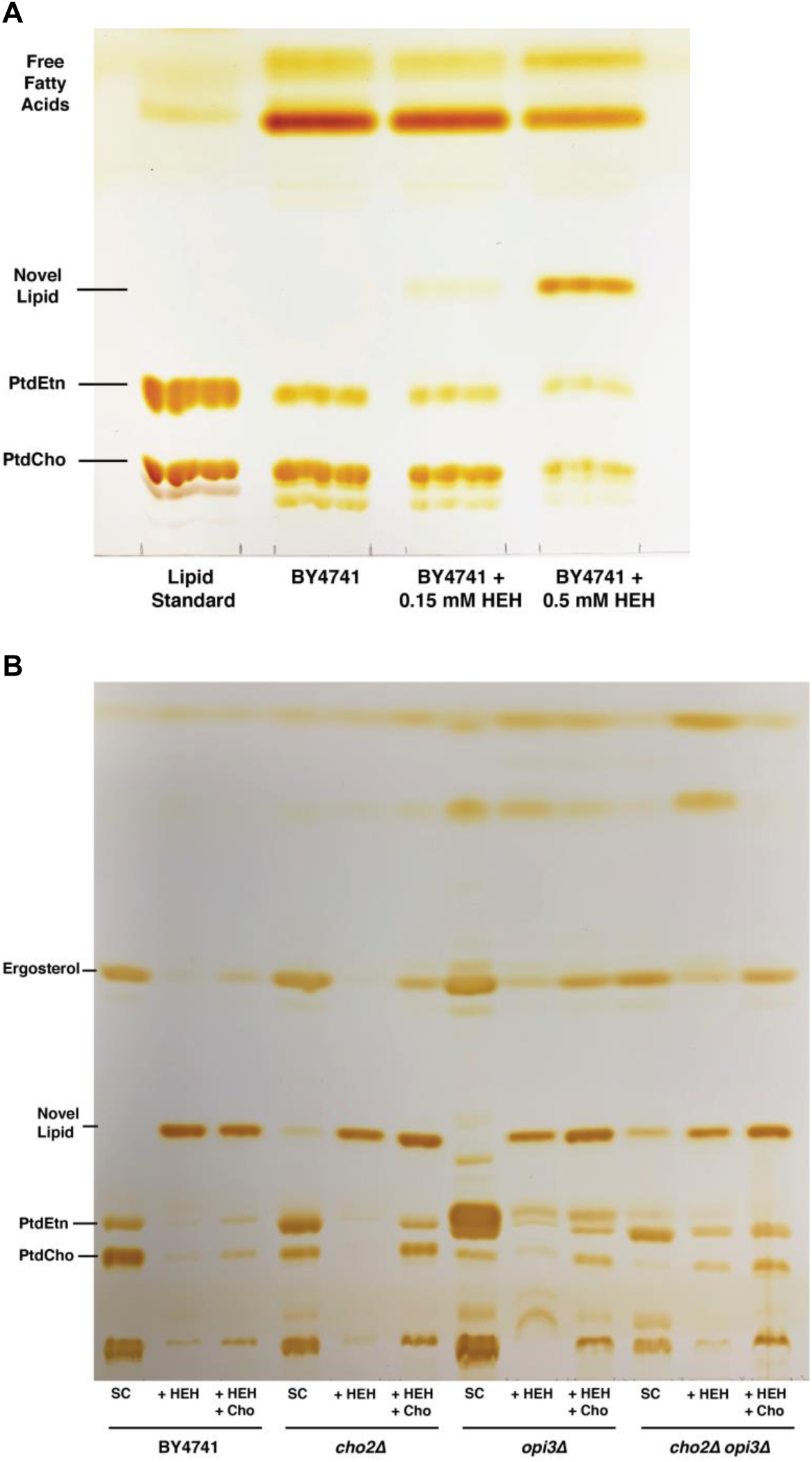
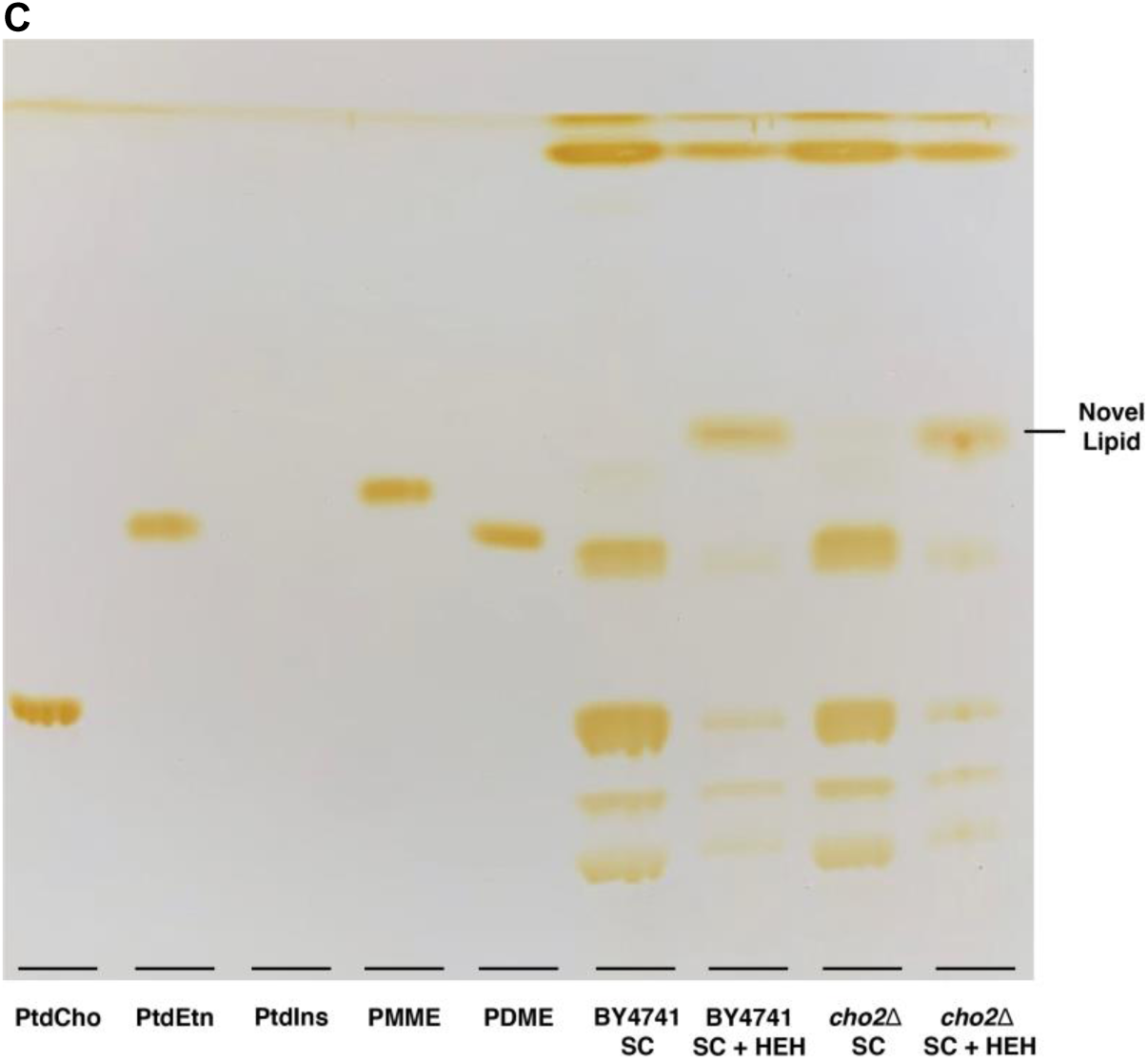
HEH supplementation induces the formation of an unidentified phospholipid. *A*, wildtype phospholipid profile following the addition of 0.15 mM or 0.5 mM HEH. *B*, WT, *cho2Δ*, *opi3Δ*, and *cho2Δopi3Δ* whole lipid profiles following supplementation of HEH or HEH and choline. *C*, comparison of WT and *cho2Δ* phospholipid profiles with known lipid standards of PtdCho, PtdEtn, PMME (phosphatidylmonomethylethanolamine), or PDME (phosphatidyl-dimethylethanolamine)

### Disruption of the choline transporter-encoding gene, *HNM1*, prevents HEH uptake and renders mutant cells resistant to HEH

As described above, Hnm1p is the principal high-affinity transporter responsible for exogenous Cho and Etnuptake (24). Due to the structural similarity of Etn and HEH, we proposed that HEH is also actively imported into the cell by the Hnm1p transporter (19). To test this hypothesis, the HEH sensitivity of the BY4741 wild-type was compared to an *hnm1Δ* strain, with or without supplementation of 0.5 mM Cho or lyso-PtdCho. The HEH dose-response curves indicated that the *hnm1Δ* strain (**Figure 3A**) is insensitive to HEH relative to the wild-type BY4741, and as expected, the presence of exogenous Cho did not alter this response to the presence of HEH. The *hnm1Δ* dose-response curve was essentially superimposable on that of the Cho-supplemented wild-type, providing strong evidence that the impairment of HEH uptake in the *hnm1Δ* mutant renders the strain resistant to the cytotoxic effects of this compound, and prevents inhibition of the PtdCho M-pathway by HEH. Supplementation of lyso-PtdCho (**Figure 3B**) very slightly increased *hnm1Δ* growth at the highest HEH concentrations, perhaps because lyso-PtdCho enters the cell through a different transport system, *hnm1Δ* is still able to access it and promote PtdCho biosynthesis through the acyltransferase pathway. Likewise, the lipid profile of *hnm1Δ* in **Figure 3C** exhibited identical contents of PtdCho and PtdEtn regardless of HEH addition, as well as the near absence of the unidentified band in both sets of samples, as described above. In the BY4741-untreated and *hnm1Δ* samples, we note the presence of a faint band in the same location as the prevalent novel band described above, which provides additional evidence that this species could be the proposed bioactive inhibitory compound, PtdHEH, or a different lipid that accumulates following HEH inhibition of the methylation pathway. Taken together, these data show that: 1. HEH enters the cell via the Hnm1p transporter; 2. Cho and lyso-PtdCho are both individually sufficient to support growth in the HEH inhibited state, and; 3. HEH inhibition leads to chemically-induced Cho or Lyso-PtdCho auxotrophy, and that this chemically induced Cho or lyso-PtdCho auxotrophy could be exploited in chemical-genomic screens for mutants that present growth defects when the methylation pathway of PtdCho biosynthesis is impaired, as detailed below.

**Figure 3:**
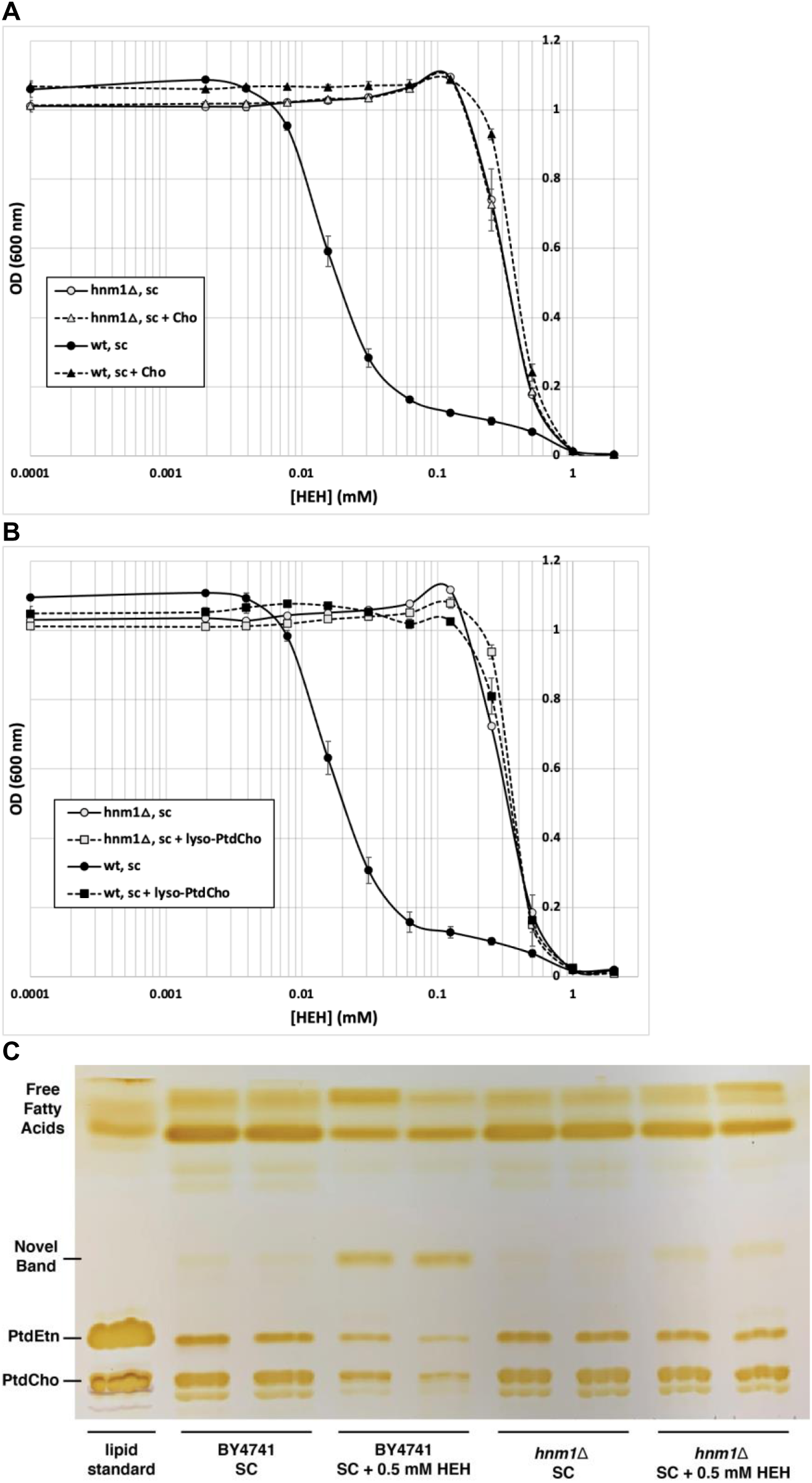
HNM1 is the HEH transporter. *A*, *hnm1Δ* growth sensitivity to HEH in synthetic complete liquid media following choline (Cho) supplementation. *B*, *hnm1Δ* growth sensitivity to HEH in synthetic complete liquid media following lyso-phosphatidylcholine (Lyso-PtdCho) supplementation. *C*, *hnm1Δ* and WT phospholipid profiles following the addition of 0.5 mM HEH, as compared to a known lipid standard containing PtdEtn and PtdCho.

### A functional genomic screen for strains that are hypersensitive to defects in the methylation pathway of PtdCho biosynthesis

To identify genes that interact epistatically with defects in the PtdCho methylation pathway, we screened a haploid *S. cerevisiae* whole-genome deletion strain collection for mutants that show enhanced sensitivity to HEH, indicated by a failure to grow in the presence of an otherwise sub-inhibitory concentration of HEH. We used a 96-pin replicator to transfer a small amount of inoculum onto an SC Glc plate, as a positive control on which all strains should grow, and an SC Glc plate containing HEH, to identify hypersensitive strains. Due to the absence of extracellular choline or lyso-PtdCho in the growth medium, the mutants were forced to rely solely on the methylation pathway for all PtdCho biosynthesis. The concentration of HEH used to isolate the sensitive mutants was determined empirically by titrating the dose needed to inhibit growth of the wild type on solid media. Concentrations >600 µM were inhibitory to the wild type, but 400 µM HEH in solid media gave robust growth of strain BY4741, and thus was used as the primary screening concentration for identification of sensitive mutants. This is slightly higher than the minimum inhibitory concentration of HEH for the wild type strain grown in liquid media, a phenomenon which has been observed in other contexts with other agents, e.g. for the cytotoxic lyso-PtdCho analog Miltefosine (25). The overall screening approach, and a typical primary screening result, are shown in **Figures 4A-B**. The mutant strains indicated by circles exhibited a complete loss of growth relative to the control plate, and all strains that showed this phenotype were identified in this primary screen. We identified ∼410 strains out of ∼4800 total, of which 371 encoded functional genes, and 39 encoded “dubious” genes, which often are adjacent to or overlapping with a different verified gene. To determine the general categories of gene function and protein localization, a Gene Ontology (GO) enrichment analysis was conducted, presented in **Table 1**. The list was characterized based on the most prevalent GO processes or components following perturbation of the PtdCHho methylation pathway, the number of genes that fell within each GO identifier term, and the associated *P*-value. Interestingly, an enrichment in mitochondrial components and processes, vacuolar acidification, and ribosomal, cytoplasmic, and membrane-bound organelle components were observed.

**Figure 4:**
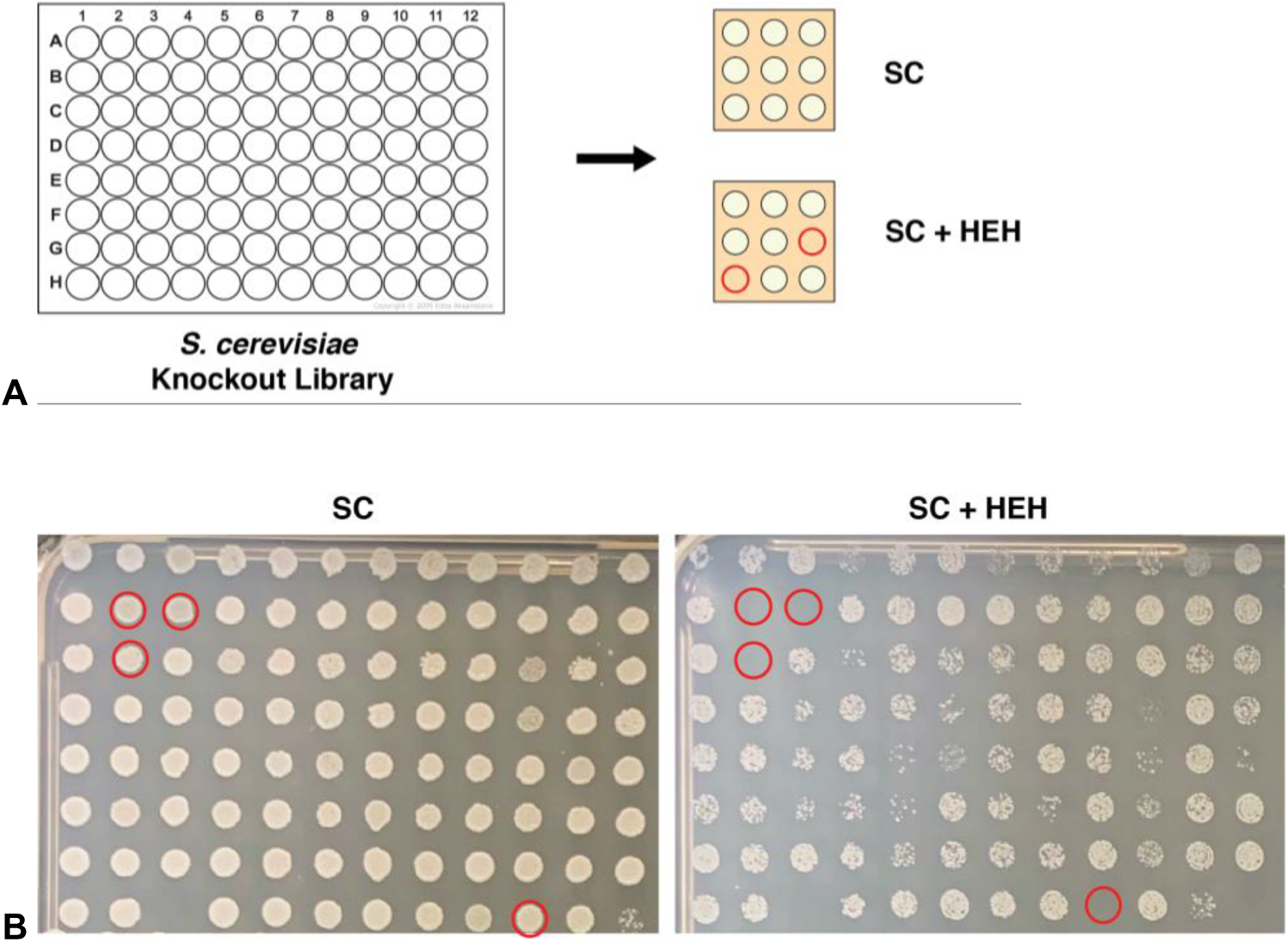
Screening schematic and results of typical sensitive mutants. *A*, visual schematic of the mutant screen. The *S. cerevisiae* MATα single-gene knockout library was inoculated onto large Petri plates containing 0- or 400 µM HEH, using a 96-pin multi-blot applicator. *B*, example screening results of a library plate and phenotype of sensitive mutants, indicated by the red circles.

**Table 1:** Gene-ontology term enrichment for preliminary gene list. The preliminary genes identified in the mutant screen were analyzed using the Gene Ontology Term Finder resource available at www.yeastgenome.org. The ontology function and component aspects were analyzed, with P < 0.001 as calculated using the hypergeometric test.

| GO Identifier | GO component (C) or Process (P) | Total for term | P-value |
| --- | --- | --- | --- |
| GO:0005759 | C: mitochondrial matrix | 66 (16.1%) | 1.11e-26 |
| GO:0140053 | P: mitochondrial gene expression | 57 (13.9%) | 3.55e-22 |
| GO:0005739 | C: mitochondrion | 154 (37.6%) | 2.31 e-21 |
| GO:0032543 | P: mitochondrial translation | 51 (12.4%) | 1.23e-20 |
| GO:0000313 | C: organellar ribosome | 37 (9.0%) | 1.63e-20 |
| GO:0044444 | C: cytoplasmic part | 284 (69.3%) | 1.90e-12 |
| GO:0016469 | C: proton-transporting two-sector ATPase complex | 18 (4.4%) | 4.17e-12 |
| GO:0005840 | C: ribosome | 50 (12.2%) | 7.24e-12 |
| GO:0007035 | P: vacuolar acidification | 17 (4.1%) | 8.57e-12 |
| GO:0043227 | C: membrane-bounded organelle | 328 (80.0%) | 6.15e-11 |
| GO:0043231 | C: intracellular membrane-bounded organelle | 319 (77.8%) | 6.04e-09 |
| GO:0015934 | C: large ribosomal subunit | 31 (7.6%) | 1.87e-08 |
| GO:0005737 | C: cytoplasm | 331 (80.7%) | 3.12e-08 |
| GO:0007005 | P: mitochondrion organization | 47 (11.5%) | 5.94e-08 |
| GO:0005740 | C: mitochondrial envelope | 61 (14.9%) | 9.52e-08 |
| GO:0006518 | P: peptide metabolic process | 88 (21.5%) | 1.67e-06 |
| GO:0000314 | C: organellar small ribosomal subunit | 14 (3.4%) | 1.74e-06 |
| GO:0031966 | C: mitochondrial membrane | 53 (12.9%) | 6.71e-06 |
| GO:0098796 | C: membrane protein complex | 38 (9.3%) | 1.19e-05 |
| GO:0006412 | P: translation | 81 (19.8%) | 4.36e-05 |
| GO:0009059 | P: macromolecule biosynthetic process | 155 (37.8%) | 4.65e-05 |
| GO:0043233 | C: organelle lumen | 108 (26.3%) | 4.69e-05 |
| GO:1904669 | P: ATP export | 9 (2.2%) | 0.00012 |
| GO:0031967 | C: organelle envelope | 65 (15.9%) | 0.00020 |
| GO:0036452 | C: ESCRT complex | 8 (2.0%) | 0.00027 |
| GO:0017004 | P: cytochrome complex assembly | 13 (3.2%) | 0.00042 |
| GO:0015992 | P: proton transport | 20 (4.9%) | 0.00063 |

Due to the large number of genes identified, we proposed that many of these strains were perhaps partially defective in growth in the uninhibited condition, and that growth inhibition due to the HEH treatment was not necessarily specific to inhibition of the methylation pathway of PtdCho synthesis. We therefore conducted more rigorous analyses of a subset of strains in order to test whether the phenotype was due to interaction with defects in the M-pathway, and were not selected. We carried out this task by analyzing the entire preliminary gene list, and identifying those genes with known genetic interactions with *cho2Δ/pem1Δ* or *opi3Δ/pem2Δ*. This process identified the genes which, when disrupted, produce a synthetic-lethal phenotype when also combined with the disruption of either cho2Δ/pem1Δ or opi3Δ/pem2Δ, suggesting that the genes are involved in processes that become indispensable when the PtdCho M-pathway is no partially inhibited by HEH. We thus identified a subset of 21 genes based on those that met all three criteria, as depicted in **Figure 5A**: 1. HEH sensitivity, and; 2. negative genetic interactions with *cho2Δ/pem1*Δ, and; 3. negative genetic interactions with *opi3Δ/pem2Δ*. The subset of 21 genes meeting these criteria are listed in **Table 2**. Manual annotation of these genes revealed that most were involved in membrane-related functions, chromatin remodeling and transcriptional regulation, and in components of the high-osmolarity/glycerol pathway.

**Figure 5:**
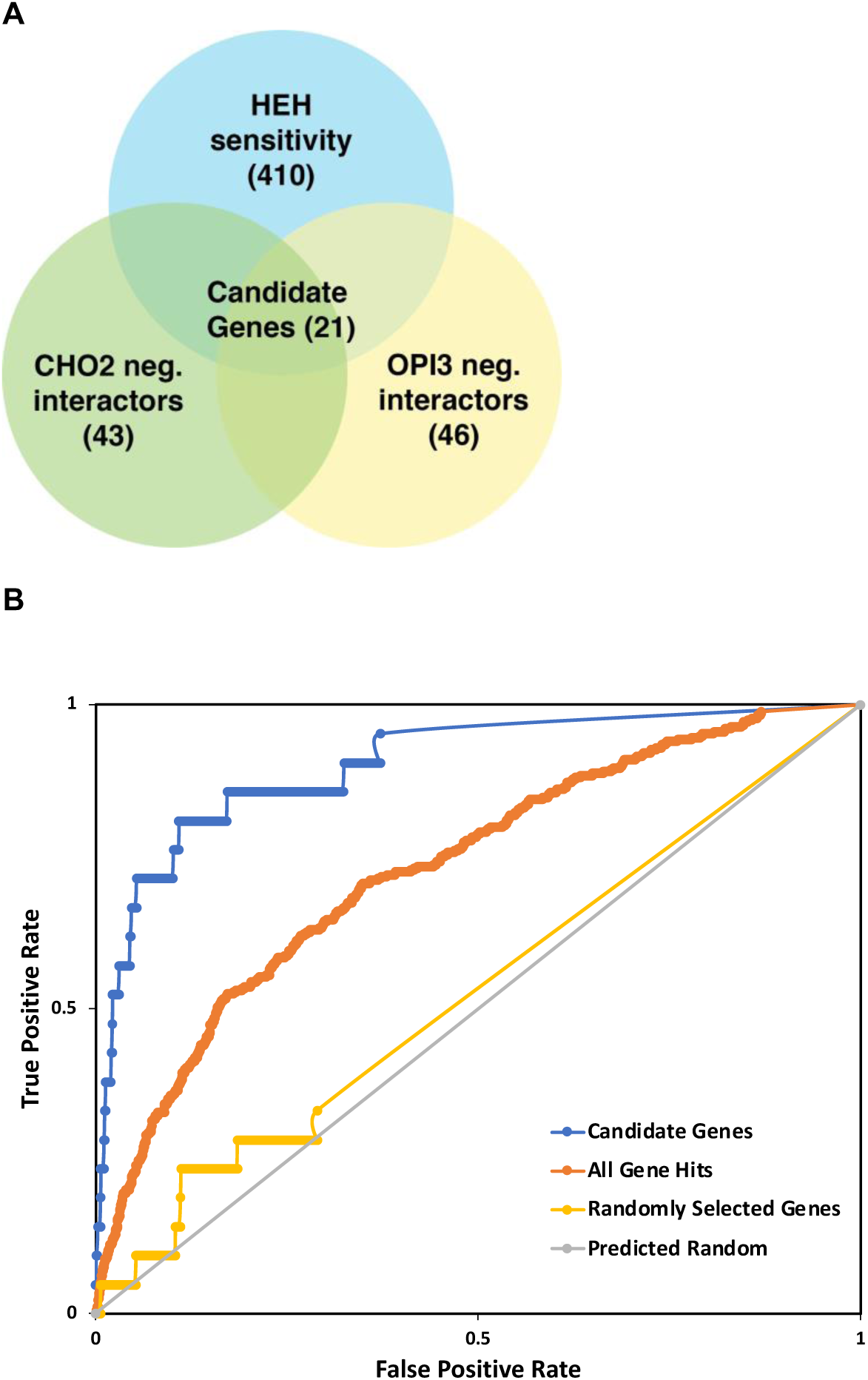
Isolation of candidate genes based on PEMT activity. *A*, criteria used to isolate the candidate genes from the preliminary mutant screen gene hits. The isolated gene set exhibited (1) sensitivity to HEH, as identified in the screen, and established negative genetic interactions with (2) CHO2 and (3) OPI3, identified via the Saccharomyces Genome Database (SGD). *B*, Receiver-operating characteristic (ROC) curve analysis of candidate gene set and preliminary gene set. Using the YeastNet gene-set enrichment analysis tool, both gene lists and a randomly selected set of genes were analyzed for level of gene connectivity and relatedness.

**Table 2:** List of candidate genes identified to interact with the methylation pathway. Gene names are listed in alphabetical order and include relative chromosomal position according to the standard SGD identifier, a description of function, and the gene localization.

| Gene ID | Gene Name | Description | Localization |
| --- | --- | --- | --- |
| YOR141C | ARP8 | Actin-Related Protein | Nucleus |
| YDR204W | COQ4 | COenzyme Q | Mitochondrion; mitochondrial inner membrane |
| YKL139W | CTK1 | Carboxy-Terminal domain Kinase | Nucleus |
| YKL213C | DOA1 | Degradation Of Alpha | Nucleus, cytoplasm |
| YNL080C | EOS1 | ER-localized and Oxidants Sensitive | Endoplasmic reticulum (membrane) |
| YLR056W | ERG3 | ERGosterol biosynthesis | Endoplasmic reticulum (lumen) |
| YML008C | ERG6 | ERGosterol biosynthesis | Mitochondrion (outer membrane), endoplasmic reticulum, lipid droplet |
| YER083C | GET2 | Guided Entry of Tail-anchored proteins | Endoplasmic reticulum |
| YJL134W | LCB3 | Long-Chain Base | Endoplasmic reticulum |
| YIR033W | MGA2 | Multicopy suppressor of GAM1 (snf2) | Endoplasmic reticulum (membrane), nucleus |
| YDR162C | NBP2 | Nap1 Binding Protein | Nucleus, cytoplasm |
| YJL117W | PHO8 | PHOspate metabolism | Endoplasmic reticulum |
| YGL167C | PMR1 | Plasma Membrane ATPase Related | Golgi membrane |
| YDL006W | PTC1 | Phosphatase type Two C | Nucleus, cytoplasm |
| YDL020C | RPN4 | Regulatory Particle Non-ATPase | Nucleus, cytoplasm |
| YDR293C | SSD1 | Suppressor of SIT4 Deletion | Cytoplasm, cellular bud, nucleus, P-body |
| YER151C | UBP3 | UBiquitin-specific Protease | Cytoplasm, cytosol |
| YEL051W | VMA8 | Vacuolar Membrane ATPase | Vacuolar V-type ATPase complex |
| YKL119C | VPH2 | Vacuolar pH | Endoplasmic reticulum (membrane) |
| YNR006W | VPS27 | Vacuolar Protein Sorting | Vacuolar membrane, endosome |
| YDR027C | VPS54 | Vacuolar Protein Sorting | Golgi apparatus, mitochondrion |

To analyze whether the mutants collected from the preliminary and candidate gene lists exhibited related functions or a diverse variety of non-shared functions, a Receiver Operating Characteristic (ROC) curve analysis was conducted using the genetic interaction and protein:protein interaction data using the Yeast Net v3 database network search for identifying new members of a pathway (26, 27). Both gene sets were compared for connectedness against a random subset of genes, with the area under the curve (AUC) value being directly proportional to the level of connectivity. As depicted in **Figure 5B**, both gene sets exhibit a high level of connectedness in terms of genetic interactions and membership in protein complexes. The narrowed set of 21 genes displayed a significantly higher degree of connectivity, with an AUC value of 0.90201 and *P*-value of 1.110335e-^12^. This is as compared to the larger gene set identified in the primary screen (AUC value 0.7326, *P*-value of 1.372305e-^36^). In contrast, a randomly selected gene list included an almost equivalent value of true and false connections, resulting in a lower AUC value (0.5292) that is, as expected, almost superimposable to and statistically indistinguishable from the predicted random association.

### Choline and lyso-phosphatidylcholine can maintain cellular phosphatidylcholine levels when the methylation pathway is chemically inhibited by HEH

Due to the redundant nature of PtdCho biosynthesis, i.e. the presence of several different pathways to the same essential molecule, sufficient PtdCho can be produced to support membrane biogenesis and cell growth in standard laboratory conditions, as long as one of the three biosynthetic pathways is functional (7). For HEH to act as a specific inhibitor of Cho2p and Opi3p, cellular HEH-mediated sensitivity should dramatically lessen following the supplementation of exogenous choline or lyso-PtdCho. In this way, stimulation of either of the two alternative PtdCho pathways (Kennedy or Acyltransferase) should overcome the effects of the single disrupted methylation pathway. We conducted a series of assays to quantify susceptibility to 2-HEH at different concentrations, as well as to qualitatively observe choline and lyso-PtdCho-derived growth recovery following PtdCho methylation pathway inhibition. In addition to using the WT strain BY4741 as a control, we also tested *hnm1Δ* and three mutant strains from the list of 21 genes: *ARP8*, which encodes a nuclear-localized actin related protein involved in chromatin remodelling; *LCB3*, encoding encoding a sphingoid-base phosphate-phosphatase involved in regulating sphingolipid content and composition; and *ERG6*, encoding a C-24 sterol methyltransferase, which is a non-essential step in the ergosterol biosynthetic pathway. We also included the *hnm1Δ* mutant as an HEH resistant control.

These results (**Figure 6A**) verify that increasing concentrations of HEH in liquid media render both *arp8Δ* and *erg6Δ* more sensitive to HEH than the WT. In SC Glc, the initial inhibitory effects were exhibited at ∼8 µM for the mutants and ∼16 µM for the WT, with the addition of 0.5 mM Cho restoring all strains to HEH resistance. This result directly indicates the PtdCho M-pathway-specific targeting of HEH, as promotion of the alternative Kennedy pathway ameliorates the sensitive phenotype. As exhibited in **Figure 1C**, a similar recovery phenotype has also been observed in the WT following lyso-PtdCho supplementation, reliant instead on the A-pathway.

**Figure 6:**
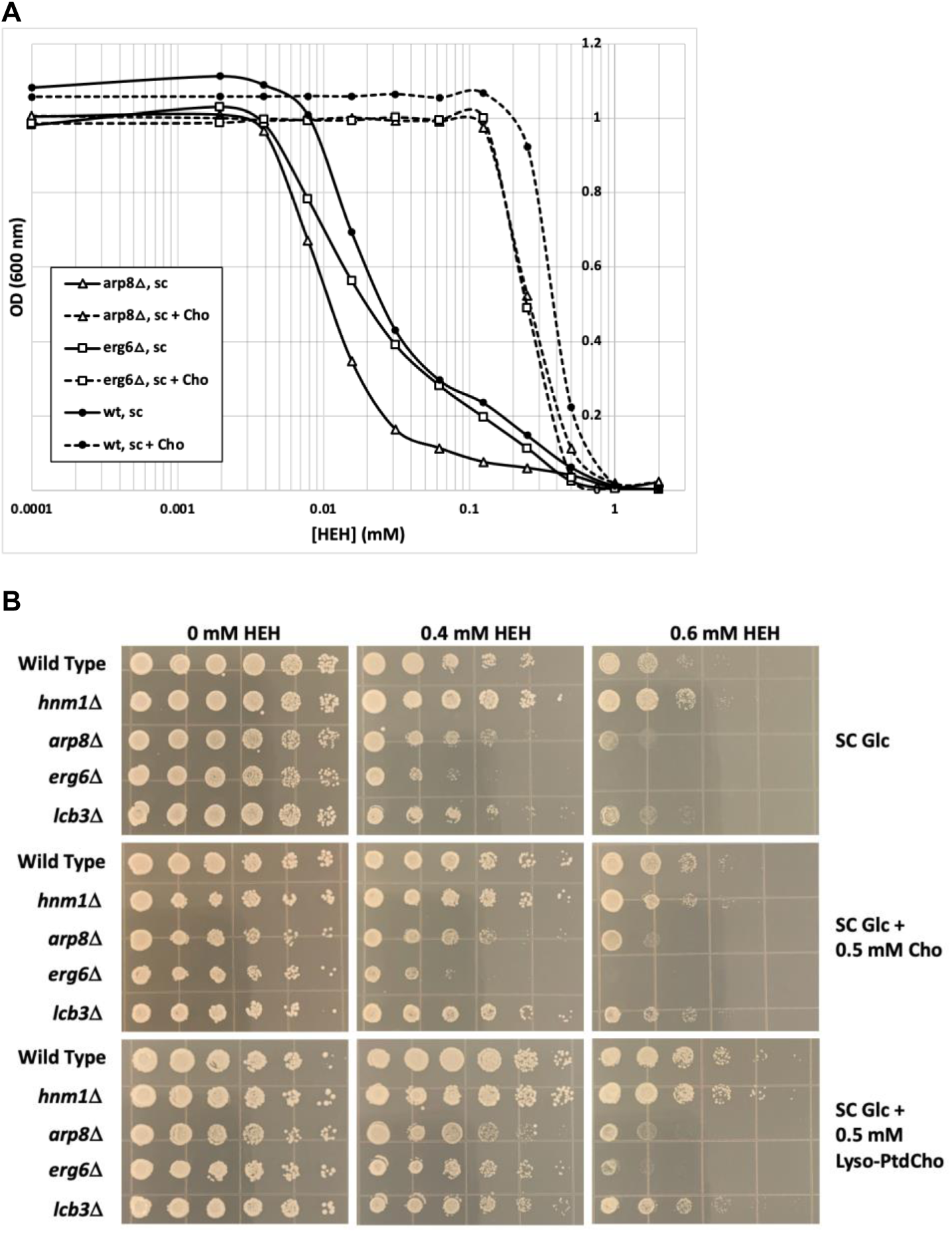
Re-screening of candidate genes. *A*, mutant growth sensitivity to HEH in synthetic complete liquid media following choline (Cho) supplementation. *B*, growth rate of WT, *hnm1Δ*, and candidate mutants following addition of HEH (0-, 0.4-, or 0.6 mM) or the supplementation of Cho or lyso-phosphatidylcholine (lyso-PtdCho). All strains were 5-fold diluted and pinned onto the respective media conditions using a 48-pin replicator.

**Figure 7:**
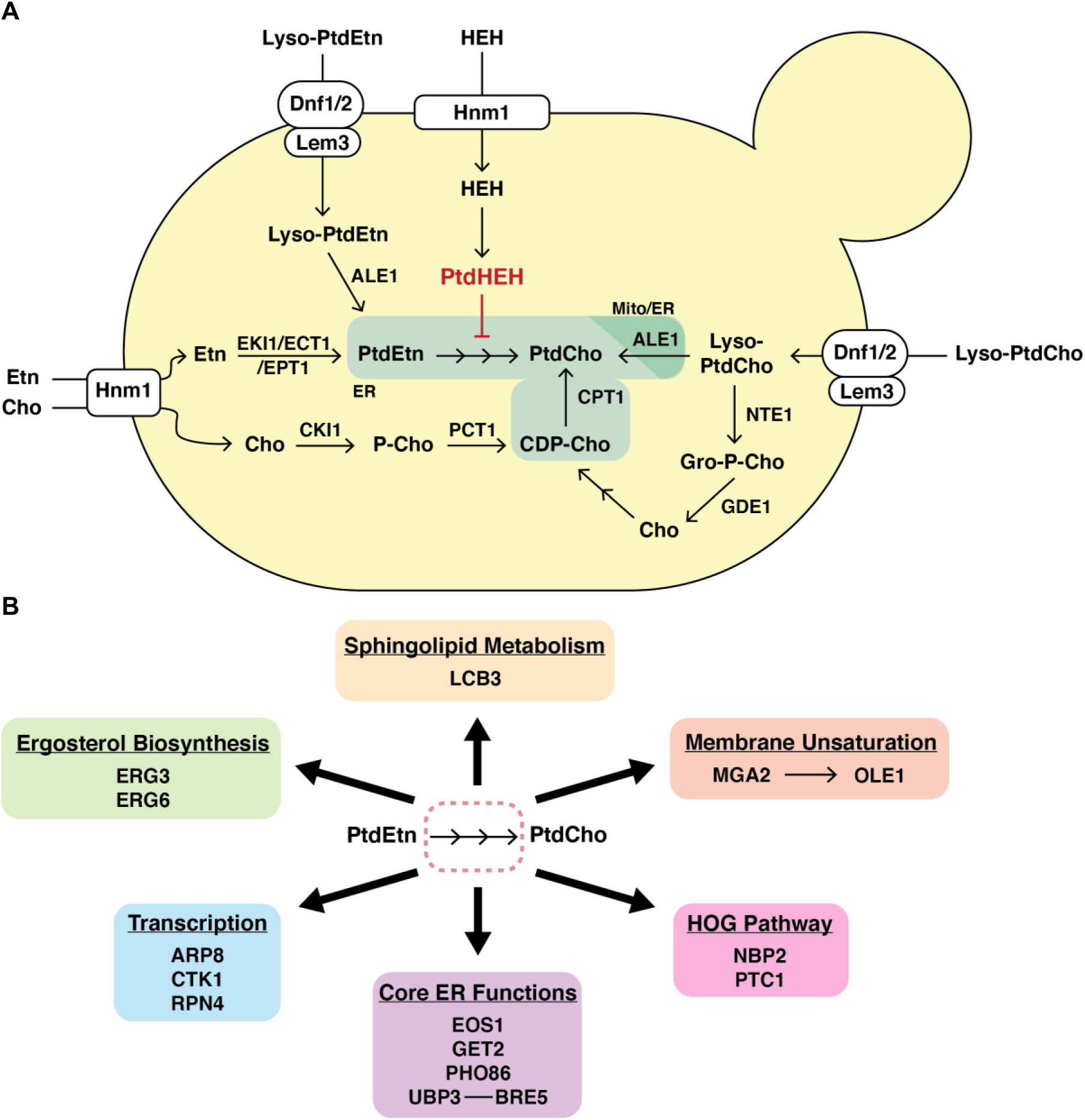
Comprehensive Model of Phosphatidylcholine biosynthesis. Comprehensive pathway diagram of the three phosphatidylcholine (PtdCho) biosynthetic pathways, with HEH-mediated inhibition of the PtdCho-methylation pathway. In this model, PtdCho can be synthesized by (1) the Kennedy pathway, in which uptake of exogenous ethanolamine (Etn) or choline (Cho) by the HNM1 transporter are converted in a series of metabolic steps into phosphatidylethanolamine (PtdEtn) or PtdCho, respectively; (2) the conversion of Etn- or Lyso-PtdEtn-based endogenously produced PtdEtn into PtdCho via the Methylation pathway’s tri-methylation of PtdEtn; or (3) the import of exogenous lyso-phosphatidylcholine (Lyso-PtdCho) by the dual action of LEM3 and DNF1/2, and subsequent acylation of Lyso-PtdCho into PtdCho via ALE1. Lyso-PtdCho can also undergo a degradation pathway, in which it is broken down into Cho and then becomes metabolized into PtdCho through the Kennedy pathway. The PtdCho-related inhibitory effects of HEH occur through its similar utilization of the HNM1 transporter, and downstream targeting of the methylation pathway in manners that remain unclear.

This analysis yielded similar results when conducted on solid media. The WT, *hnm1Δ,* and strains selected for further characterization (*arp8Δ*, *lcb3Δ*, and *erg6Δ*) all exhibited increased sensitivity when serial dilutions were pinned to solid media, similar to the conditions in the initial screen, and subjected to increasing concentrations of HEH on minimal media plates (SC Glc), as shown in **Figure 6B**. Additionally, the supplementation of Cho and lyso-PtdCho both independently reduced this sensitivity and thus promoted growth recovery, with lyso-PtdCho supporting a greater degree of recovery than Cho. This result indicates that upon disruption of the PtdCho methylation pathway, cells may better utilize A-pathway derived PtdCho to replenish cellular PtdCho pools. When compared to the WT, *hnm1Δ* grows at a similar rate in all supplemented conditions, and even outcompetes WT growth in minimal SC Glc medium, further confirming that HEH’s effects on PtdCho biosynthesis are dependent on Hnm1p-mediated cellular uptake. Within this restricted gene set for re-screening, *erg6Δ* displayed the greatest sensitivity across all conditions, which perhaps indicates that there is an essential membrane function or biophysical property, defects in which are tolerated when either ergosterol biosynthesis or the PtdCho M-pathway are partially disrupted, but not both pathways at the same time. This suggestion about the relationship between these two processes is bolstered by the observation that exogenous Cho or lyso-PtdCho failed to fully rescue HEH-inhibited *erg6Δ* cultures. Finally, *lcb3Δ* was observed to outcompete *arp8Δ* in all media conditions, especially during lyso-PtdCho addition. In a similar manner to Erg6, it is likely that the genetic function of Arp8 in chromatin remodeling has a more important relationship with the PtdCho methylation pathway than Lcb3, involved in sphingolipid metabolism.

## Discussion

This study aimed to systematically identify genes which, when disrupted, lead to a synthetic growth defect with the M-pathway of PtdCho synthesis. The M-pathway, and the genes specifying it, have been known and studied since the early 1980’s, as have the K-pathway, and the more recently described A-pathway (6). As outlined in **Fig. 1B**, most eukaryotes have the capacity to synthesize PtdCho via these three separate pathways, although different organisms and different tissues preferentially utilize certain pathways over others. Generally, in the yeast model system, as long as one of these pathways is functional, the cell can synthesize sufficient PtdCho to support growth and cellular processes. The methylation pathway remains largely unexplored beyond the primary genes involved in it, which fueled our interest in developing a greater understanding of it and its genetic interactors. The compound 2-hydroxyethylhydrazine (HEH) was identified (19) as a specific PtdCho M-pathway inhibitor, however this singular study left many questions unanswered regarding the mechanistic function of HEH (19). As depicted in **Figure 1B**, the mechanistic basis for this inhibition lies in the structural similarity of 2-HEH to ethanolamine, resulting in its theorized subsequent uptake through the choline transporter Hnm1p and metabolic alterations via the Kennedy pathway.

If cellular uptake of HEH, and subsequent formation of PtdHEH (**Fig. 1A-B**), results in a direct inhibition in PtdCho biosynthesis, then a lipid profile of 2-HEH-treated cells should reflect a decrease in PtdCho levels. This result was confirmed in **Fig. 2A**, in which the concentrations of cellular PtdCho and PtdEtn were both observed to decrease, more significantly in the former. An unexpected result was the formation of a novel lipid band solely in the HEH-treated samples. Although we originally hypothesized its identity as being the phosphatidyl-form of HEH (PtdHEH), we determined that this lipid is not an HEH-derivative due to the production of this lipid (even in the absence of HEH) in the *cho2Δ*, *opi3Δ*, and *cho2Δ opi3Δ* mutants. We further validated that this species is not the PtdEtn-methylation intermediates PMME or PDME, which could have resulted from partial inhibition of the methylation pathway by HEH. The identity of this lipid remains unknown, and our results indicate that it is an endogenously synthesized lipid that accumulates upon disruption of the PtdCho methylation pathway.

We were also able to validate the role of the Hnm1p transporter in cellular uptake of HEH. The lipid profile of the disrupted HNM1 transporter (*hnm1*Δ) produced an identical phenotype to the WT-untreated sample, in both the presence and absence of HEH, as well as the distinct absence of the novel band (**Fig. 3C**). Additionally, *hnm1*Δ growth at increasing concentrations of HEH was comparable to the choline- and lyso-PtdCho supplemented growth recovery of HEH in the WT (**Fig. 3A-B**). Finally, the growth of *hnm1*Δ on solid media in **Fig. 6B** was comparable and slightly exceeded WT growth patterns when supplemented with choline and lyso-PtdCho at increasing HEH concentrations. Because choline and HEH both enter the cell through the same transporter, Hnm1p, we were curious if the choline-mediated growth rescue was due in part to a competition for entry between the two compounds. However, the similarity in recovery pattern between lyso-PtdCho and choline indicate that this is not a concern. These results directly exhibited the necessity for proper Hnm1p functioning in order to take up HEH and therefore undergo inhibited PtdEtn methylation. Our results collectively corroborate the role of Hnm1p in the transport of 2-HEH from the external to the intracellular environment.

To genetically dissect the methylation pathway for potential interactors, we designed and conducted a mutant screen to identify any nonessential genes that, when deleted, render the resulting strain unable to grow when entirely reliant on the methylation pathway for all PtdCho biosynthesis. When challenged with a concentration of HEH (400 mM) that permitted robust wild type growth, we identified 410 hypersensitive mutants with defects in non-essential genes. Of those, 371 mutants encoded functional genes, with the remainder accounting for “dubious” genes, which often overlap functional genes, and thus phenocopy a loss of function in the bona fide overlapping gene. While some mutants exhibited a slow growth phenotype upon HEH challenge, we focused on more detailed characterization of strains that showed a a complete loss of growth. Following a Gene Ontology (GO) analysis of the preliminary gene list with a focus on biological processes, the most significant GO categories included mitochondrial gene expression & translation, vacuolar acidification, mitochondrion organization, & peptide metabolic process (**Table 1**).

Additional bioinformatic analyses (**Fig. 5A**) led to the identification of 21 genes, listed in **Table 2**, which showed a negative genetic interaction with both cho2Δ and opi3Δ individually, and which were hypersensitive to HEH. A substantial majority of these genes were identified as having functions relating to membrane function, and a few were shown to have nuclear functions in chromatin remodeling and transcriptional regulation. As seen in **Fig. 5B**, the restricted gene set showed a significant level of network connectivity, as indicated by a Receiver Operating Characteristic AUC value of close to 1.0. This was not a terribly surprising result, as each of the members of this narrowed gene list were selected or interactions with PtdEtn methyltransferase defective mutants, and thus should display a high level of associated genetic and biochemical functional interactions. That said, the full mutant list of 410 genes displayed a very high AUC value, suggesting a profound overrepresentation of network connectivity relative to that expected by random chance. This result confirms that, in addition to the narrowed list of candidate genes, the entire mutant screen gene set is, individually, more likely to interact genetically or biochemically with other genes on the list than with a randomly selected subset of genes, indicating that the screen has identified a genetically and biochemically tractable subset of the yeast genome/proteome.

In summary, the data presented in this study provides new insights into the function and regulation of the methylation pathway of phosphatidylcholine biosynthesis, and extends our understanding of HEH and its function as a potent and specific PtdEtn methyltransferase inhibitor. HEH is thus an effective chemical tool that will be used to further probe and dissect PtdCho biosynthetic pathways in *S. cerevisiae* and other eukaryotic systems. This chemical-genetic screen has thus identified previously unknown genetic and biochemical interactions of a broad subset of the yeast genetic repertoire, and mechanistic biochemical elucidation of the nature of these interactions will

## Experimental procedures

### Strains and growth conditions

The haploid non-essential gene deletion collection in the BY4742 background (Mat α collection; 21) was purchased from TransOMIC Technologies (Huntsville, AL), and stored in glycerol-containing 96-well plates at -80°C. 2-hydroxyethylhydrazine (HEH), choline chloride, ethanolamine hydrochloride, and Tergitol NP-40 were purchased from Sigma, oleoyl-lyso-PtdCho was purchased from Avanti Polar Lipids, and all components for standard yeast media were purchased from US Biologicals. Routine strain growth was conducted on YPD (1% w/v yeast extract, 2% w/v peptone, 2% w/v glucose) with 2.0% w/v agar. Media used for screening consisted of SC glucose medium (1% sodium chloride, 0.34% yeast nitrogen base without amino acids, 10% 10x amino acids, 10% 10x glucose) with 2.0% w/v agar. HEH was purchased as a 1 M stock solution and diluted as necessary. Choline was prepared as a 1 M stock solution, stored at 4°C, and diluted as needed. Lyso-Phosphatidylcholine was prepared as a 10 mM stock solution in 10% filter-sterilized Tergitol, and stored at -20°C.

### Genetic Screen

All mutants in the non-essential gene deletion collection were diluted into 100 mL of SC media in a sterile 96-well plate, using a 96-pin replicator (Dan-Kar corporation, model MC 96). The diluted cultures then were inoculated onto solid media conditions in 20 by 20 cm square Petri plates using the 96-pin replicator. Media used for screening consisted of YPD and SC Glc containing 0-, 0.4-, and 0.8-mM HEH. Following inoculation, the screen plates were photographed and scored for growth following a 2–3 day incubation at 30°C. To sterilize the replicator between dilution plates, the pins were submerged in a water and bleach bath followed by ethanol flame sterilization.

### Bioinformatic analysis

The genetic screen led to the identification of 373 genes that when disrupted, experienced a complete loss of growth upon addition of 2-HEH. Using the Saccharomyces Genome Database website (SGD; www.yeastgenome.org), all gene hits were analyzed for gene function and localization and then categorized according to general function. The genetic interactions tool on the SGD allowed for all gene hits to be examined for observed interactions with osh2/pem1 and opi3/pem2; the genes that met all 3 criteria (2-HEH sensitivity and known pem1 <u>and</u> pem2 interactions) were isolated into a set of 21 candidate genes that were further analyzed moving forward.

### Analysis of WT & hnm1Δ lipid composition

Strains were grown overnight in 50 mL of SC media at 30 °C, diluted in 500 mL flasks to a 50-mL culture with an optical density (600 nm) of 0.1 per 100 mL of culture using a plate reader, and then incubated at 30 °C for 2 hrs until the optical density reached 0.2 per 100 µL of culture. Then, various concentrations of 2-HEH were added to their own culture flask (0 mM control, 0.15 mM, or 0.5 mM 2-HEH) and the flasks were incubated at 30 °C for 24 hours after the timepoint that 2-HEH was added. 10-mL aliquots were then removed from each flask, washed 2x with 1x PBS, then the supernatant was removed. The samples were stored in the -20 °C freezer.

Lipid extracts were prepared from cell pellets by an ethanolic-Bligh-Dyer lipid extraction (22) as described in Riekhof and Voelker, 2006. The chloroform phase was removed with a Pasteur pipette and dried using a nitrogen evaporator (N-EVAP^TM^ 112 with OA-SYS Heating System, Organomation Assoc., Inc.). Lipids were stored as dried films at -20 °C prior to analysis. The lipid extracts were suspended in 200 mL of chloroform/methanol (9:1, v/v), of which 50 mL was spotted onto the TLC plate. A solvent system of chloroform, acetone, methanol, glacial acetic acid, and water (50:20:10:10:5) was used, followed by visualization of lipid bands in an iodine vapor filled chamber.

### Analysis of cho2Δ, opi3Δ, & cho2Δopi3Δ lipid composition

All strains, including the WT, were grown overnight in 25 mL of YPD liquid media at 30 °C, washed 2x in fresh SC Glc, and resuspended in 10 mL SC Glc. Washed cells were diluted to 0.1 OD in 50-mL cultures in SC Glc, then incubated at 30 °C until the OD reached 0.2 per 100 µL of culture. Cultures were supplemented with the following conditions: 0.4 mM HEH, 0.5 mM choline, 0.4 mM HEH + 0.5 mM choline, or a no supplement control. After a 24-hour growth period following the supplement addition, cultures were spun down, washed 2x with 1x PBS, removed the supernatant & stored the samples at -20 °C. The samples then underwent a whole lipid extraction protocol as outlined above and analyzed for phospholipid content via TLC.

Comparison of the WT and *cho2Δ* phospholipid profiles to the PtdCho methylation pathway intermediates was completed using the lipid extracts obtained from this experiment, as well as PMME and PDME lipid standards obtained from Avanti Lipids. Samples were spotted onto a TLC plate and analyzed using the same solvent system mentioned above, followed by iodine chamber-mediated visualization.

### HEH Susceptibility Assays

All growth assays were conducted in 96-well plates in which HEH was used in a 2-fold dilution series in SC Glc media. The final concentrations of HEH ranged from 2 mM to 0.001953125 mM, with a 0 mM negative control. All strains were grown overnight in 5 mL of SC Glc at 30 °C, then 1:200 diluted into fresh SC Glc or SC Glc supplemented with 0.5 mM-choline (SCC), ethanolamine (SCE), or lyso-PtdCho (SCL), and then added to the 96-well dilution plates. Dilution series were assessed in quadruplicate and included the WT strain BY4741, the parental strain for the mutant collection. Plates were incubated at 30 °C for 24 hrs, then growth was measured as optical density at 600 nm on a plate reader. For analyses on solid media, strains were grown overnight in 5 mL of SC Glc at 30 °C, then diluted as a sequential 5-fold series in SC Glc media in the wells of a 96-well plate. A 48-pin replicator tool was used to transfer ∼2 µl aliquots of the culture dilutions onto SC Glc, SCC, or SCL agar plates that contained 0-, 0.4-, or 0.6-mM HEH. Following inoculation, the plates were photographed and scored for growth following a 2–3 day incubation at 30°C.

## Data Availability

All data used to generate figures, or otherwise described in the text, are available from the corresponding author

## Author Contributions

AM designed and conducted experiments and wrote the manuscript, WR designed and coordinated experiments and edited the manuscript draft.

## Conflict of Interest

None declared.

## Abbreviations

HEH: 2-Hydroxyethylhydrazine
PtdCho: phosphatidylcholine
PtdEtn: phosphatidylethanolamine
PMME: phosphatidylmonomethylethanolamine
PDME: phosphatidyldimethylethanolamine
Cho: choline
Etn: ethanolamine
lyso-PtdCho: lyso-phosphatidylcholine
SGD: Saccharomyces genome database
ROC: receiver operating characteristic
IC50: half maximal inhibitory concentration
MIC: minimal inhibitory concentration

## Notes

### Competing Interest Statement

The authors have declared no competing interest.

